# Application of Airy beam Light sheet microscopy to examine early neurodevelopmental structures in 3D hiPSC-derived human cortical spheroids

**DOI:** 10.1101/2020.06.27.174904

**Authors:** Dwaipayan Adhya, George Chennell, James Crowe, Eva P. Valencia-Alarcón, James Seyforth, Neveen Honsy, Marina V. Yasvoina, Robert Forster, Simon Baron-Cohen, Anthony C. Vernon, Deepak. P. Sriavstava

**Affiliations:** Department of Basic and Clinical Neuroscience, Maurice Wohl Clinical Neuroscience Institute, Institute of Psychiatry, Psychology and Neuroscience, King’s College London, London, UK; Autism Research Centre, Department of Psychiatry, University of Cambridge, Cambridge, UK; MRC Centre for Neurodevelopmental Disorders, King’s College London, London, UK; M Squared Life Ltd., The Surrey Technology Centre, 40 Occam Road, Guildford, UK; Department of Neuroimaging, Institute of Psychiatry, Psychology and Neuroscience, King’s College London, London, UK

## Abstract

**Background:** The inability to observe relevant biological processes *in vivo* significantly restricts human neurodevelopmental research. Advances in appropriate *in vitro* model systems, including patient-specific human brain organoids and human Cortical Spheroids (hCSs) offer a pragmatic solution to this issue. In particular, hCSs are an accessible method of generating homogenous organoids of dorsal telencephalic fate, which recapitulate key aspects of human corticogenesis, including the formation of neural rosettes. These neurogeneic niches give rise to neural progenitors that subsequently differentiate into neurons. Atypical formation of these structures has been associated with neurodevelopmental disorders such as autism spectrum conditions, from studies of patient-specific human induced pluripotent stem cells grown as 2D cultures. Thus far however, conventional methods of tissue preparation in this field limit the ability to image these structures in three-dimensions within intact hSC or other 3D preparations. To overcome this limitation, we have sought to optimise a methodological approach to process hCSs to maximise the utility of a novel Airy-beam light sheet microscope (ALSM) to acquire high resolution volumetric images of internal structures within hCS representative of early developmental time points.

**Results:** Conventional approaches to imaging hCS by confocal microscopy were limited in their ability to image effectively into intact spheroids. Conversely, volumetric acquisition by ALSM offered superior imaging through intact, non-clarified, *in vitro* tissues, in both speed and resolution as compared to conventional confocal imaging systems. Furthermore, optimised immunohistochemistry and optical clearing of hCSs afforded improved imaging at depth. This permitted visualization of the morphology of the inner lumen of neural rosettes.

**Conclusion:** We present an optimized methodology that takes advantage of an ALSM system that can rapidly image intact 3D brain organoids at high resolution while retaining a large field of view. This imaging modality can be applied to both non-cleared and cleared *in vitro* human brain spheroids derived from hiPSCs for precise examination of their internal 3D structures. Furthermore, this process represents a rapid, highly efficient method to examine and quantify in 3D the formation of key structures required for the coordination of neurodevelopmental processes in both health and disease states. We posit that this approach would facilitate investigation of human neurodevelopmental processes.

## Introduction

Human brain organoids represent a revolutionary step forward in our ability to investigate human neurodevelopment through the use of patient-specific human induced pluripotent stem cells (hiPSC) to generate neural tissues [1–3]. Brain organoids recapitulate a more native state of neurodevelopment, and thus have enabled the *in vitro* study of molecular and cellular events during early human neurodevelopment [4–7]. Early development is also emerging as a key period for the emergence of atypical cellular and molecular processes thought to contribute to the likelihood of neurodevelopmental disorders such as autism spectrum conditions (here after referred to as autism) [8–12]. Therefore, brain organoids offer a unique opportunity to study pathophysiological mechanisms thought to occur in these disorders [8, 10–13]. Several different methods have been used to generate 3D brain organoids from hiPSCs that recapitulate the complex cellular developmental processes involved in the generation of specific brain regions [1–3, 14–16]. One example is the generation of human cortical spheroids (hCSs), which generate dorsal cortical cells [2]. In this method, stem cells differentiate into neural progenitor cells (NPCs) that self-organise around a central lumen called a ‘neural rosette’ [2, 17] These neural rosettes display apical-basal polarity, and are thought to represent the formation of the neural tube during early brain development [2, 17, 18]. Thus, brain organoids like hCSs, provide a 3D *in vitro* system relevant for studying mechanisms relevant for human neurodevelopment and disorders such as autism.

At present, microscopic analysis of human brain organoids is primarily undertaken by conventional methodologies such as confocal microscopy, which require deconstruction of the 3D structure [1, 2, 15]. This approach typically involves imaging of 2D tissue slices of 3D organoids. This resulting in a loss of structural integrity and thus the phenotypic resemblance to their original tissues. Hence, the significance of understanding cellular processes, organisation and complexity within a system that mimics native human tissue in a 3D state is lost [19]. To image intact nervous tissues, high-speed, high-resolution volumetric imaging methods are required over large fields-of-view. New imaging modalities such as, light sheet microscopy (LSM) is rapidly gaining importance for imaging intact biological specimens [20, 21]. The advantage of using light sheet microscopy over confocal microscopy is that it rapidly (500x faster) produces 3D volumetric images and due to the speed produces less photo damage to the specimen. However, two important factors need to be taken into consideration when utilizing this approach: (1) due to lipid content, there is considerable scattering of light in imaging of large volumes of tissue that may result in a rapid degradation of signal with increasing depth of tissue; and (2) antibodies are often accumulate in superficial regions of tissue, thus reducing visibility of target proteins in deeper sections of the tissue.

Several methods are now available to render intact tissues optically transparent [22–25]. These aqueous clearing techniques reduce light scattering by removing tissue lipids and replacing them with a clear matrix, and as reported by Xu et al (2017) can also facilitate greater penetration of antibodies. The combined effect enables significant preservation of both tissue ultrastructure and the accessibility of native proteins to antibody probes [22, 24, 25]. To date only a limited number of studies have applied the use of LSM or tissue clearing in combination with human brain organoids. For example, Renner and colleagues used cleared cerebral organoids to visualise the organisation of ventricles and connectivity between different regions within cerebral organoids [26]. However, in this study, the authors utilized conventional imaging approaches that requires the manual reconstruction of 3D structures from 2D images, which is both time consuming and may result in the loss of the inherent complexities of the biological specimen [19]. Ina different study, Li and colleagues used LSM to examine cortical folding in cerebral organoids [27], but not to examine the organisation of cellular structures internally. To our knowledge, few if any studies have used a combination of LSM with tissue clearing to examine the organisation of internal cellular structures in human brain organoids.

In this study, we have: 1) compared conventional tissue processing and image acquisition methods (confocal microscopy) against a novel methodology of presenting intact 3D tissue for volumetric acquisition using LSM; and 2) optimised immunohistochemical approaches in combination with tissue clearing to improve acquisition of neural rosettes deep within hSCs. Specifically, we imaged intact hCSs generated from hiPSCs, using an Airy-beam light sheet microscopy (ALSM) system [21], in combination with an optimized immunhistochemical protocol based on the FACT approach [25] and a refractive index (RI)-matched fast tissue clearing method based on the ScaleS [24]. We confirm that the ALSM is superior in imaging through intact, non-clarified, *in vitro* tissues, both in speed and resolution as compared to conventional confocal imaging systems. Furthermore, optimised immunohistochemistry and clearing of hCSs allowed for enhanced visualisation of internal structures. This enabled the visualisation of the inner lumen of neural rosettes in 3D, and indicated that these structures may develop as a polarised tube-like structure. These finding indicate that ALSM imaging in combination with optimized imuunohistochemistry and tissue clearing promises to yield significantly better 3D imaging of internal structures in brain organoids, and thus will facilitate the study of neurodevelopmental processes relevant for typical and atypical development.

## Results

### Application of Airy-beam Light sheet microscopy (ALSM) with high resolution

A particular advantage of ALSM is that it has a wider field of view (FOV) due to the properties of the propagation-invariant Airy beam – the light sheet can traverse the FOV and maintain its properties [21] **(Figure 1A-C)**. This in-turn means that the high axial resolution is maintained across that FOV without the need for compromising the FOV or resolution, unlike other light sheet systems. The Airy-beam delivers a field of view 20 times larger than a Gaussian beam and 8 times larger than a Bessel beam equipped on the same system [21]. The Airy-beam also provides an axial resolution comparable to that of the Gaussian and twice as good as the Bessel beam [21]. The Airy-beam’s characteristic asymmetric excitation pattern creates lobes spreading the beam across the FOV, lowering the overall light exposure to the sample resulting in 80% less photo-bleaching in comparison to the Gaussian beam [21]. Furthermore, it only requires a single exposure per z slice compared to the multiple acquisitions required for some light sheet approaches, further reducing photo-toxicity. The ALSM system uses 2 water dipping objectives. The geometry of the objectives means the maximum sample area the light sheet can image is approximately 3.5 mm in depth (XZ axis) and 3.5 mm in width (X axis), but along the y-axis it is possible to have a dimension larger than 3.5 mm **(Figure 1A and B)**.

**Figure 1.**
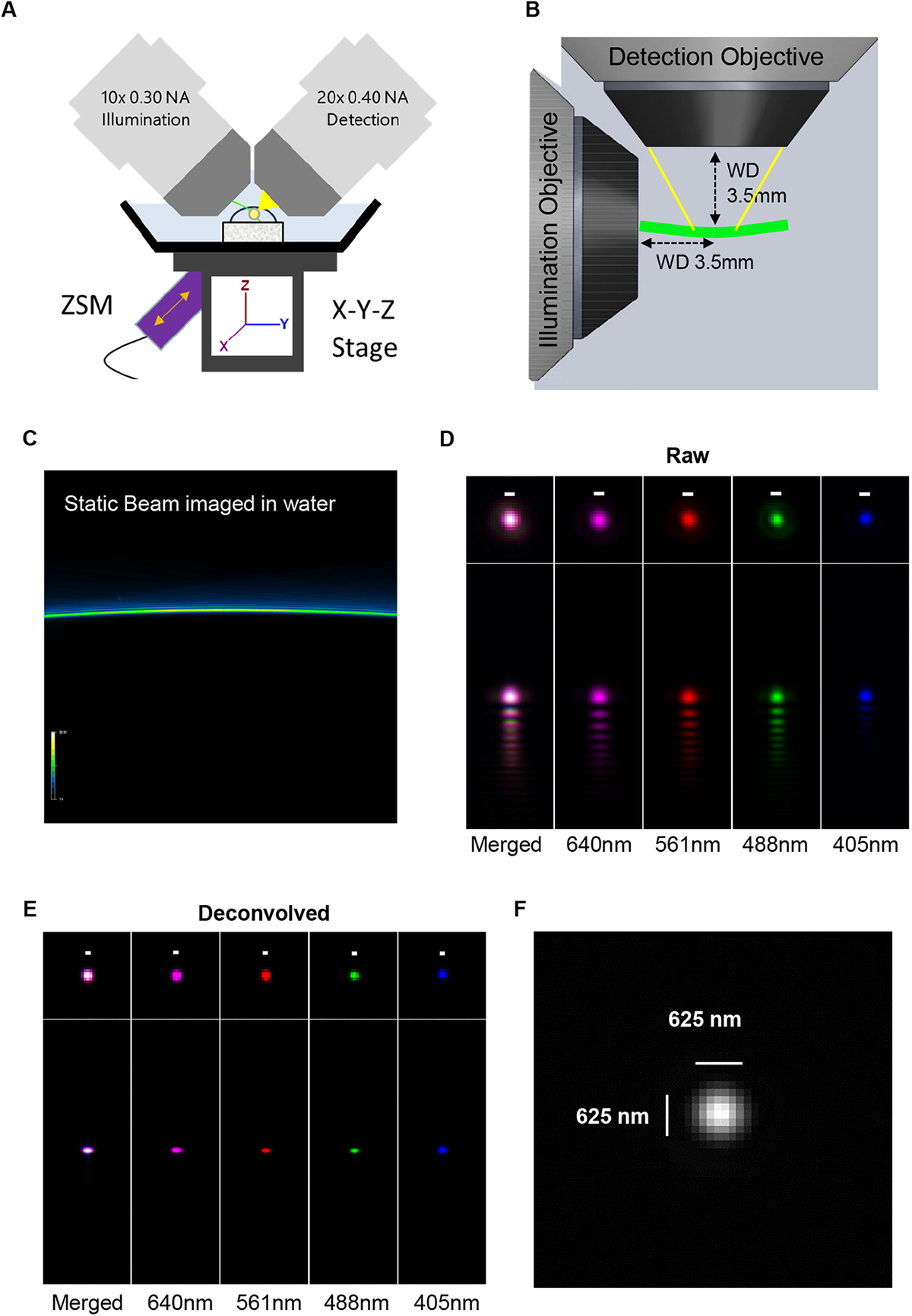
Setup and resolution of Airy-beam light sheet microscopy (ALSM). **(A)** ALSM system is equipped with 2 water dipping objectives – 10x 0.4 NA illumination and 20X 0.4 NA detection objective – positioned at a 45° angle. Biological specimens are placed on the X-Y-Z stage. Movement of the stage is controlled by a Z-axis Stepper motor (ZSM) which moves the sample along the optical axis of the detection objective. The system is equipped with 405nm, 488nm, 561nm and 640nm wavelength lasers. **(B)** Illumination objective focuses the Airy-beam (green) on to sample and the detection objective captures the emission (yellow) at 90 degrees. The geometry of the objectives gives the system a working distance of 3.5mm. **(C)** An example of the Airy-beam produced by focusing the 488nm laser though water. **(D)** Representative image of a 500 nm Tetraspeck fluorescent beads imaged at 405, 488, 561 and 640 wavelengths; image has not undergone deconvolution. Characteristic Airy lobes can be seen in the axial plane. **(E)** Deconvolved image of the fluorescent bead in C). Axial Airy lobes have been removed following deconvolution. **(F)** Image of a single 500 nm Tetraspeck fluorescent bead; the PSF of a single 500 nm bead is 0.625 μm diameter at full width at half maximum (FWHM) when imaged at 488 nm. Scale bars = 500 nm unless stated otherwise.

The resolution of the ALSM system used in this study was determined by measuring 500 nm sub-resolution Tetraspeck fluorescent beads at multiple wavelengths. Acquired raw images of beads resulted produced data stack that upon reslicing Airy lobes were visible in the axial plane **(Figure 1D)**. All bead stacks were deconvolved [21] using an extracted bead PSF and processed using Richardson-Lucy deconvolution (MSquared Cubes software). This restored the image, increasing the signal-to-noise and removed the geometric curvature of the Airy-beam **(Figure 1E)**. Measurements of resolution was assessed by determining the “Full width at half maximum” (FWHM) at the 488 nm wavelength of resliced beads where the mean system has a resolution of 0.625 μm **(Figure 1F)**.

### Challenges associated with imaging of hCSs using confocal microscopy

In order for us to assess the utility of ALSM for the imaging of 3D organoid cultures, we differentiated 3 well characterised hiPSCs [8, 28] using a free-floating directed culture method to generate 3D cortical spheroids (hCSs) [2] **(Supplemental Figure 1A-E**). At day 7, hCS specimens were ~100-300 μm in size **(Supplemental Figure 1C).** Cryo-sectioning of day 7 hCS often resulted in tissue damage. Therefore, day 7 hCSs and imaged as ‘wholemount’ intact samples by conventional confocal microscopy. As expected hCS generated from hiPSCs demonstrated the key hallmarks of *in vitro* cortical differentiation [2, 6, 17]. At this time point, hCSs were positive for the early neuroepithilium markers Nestin and Sox2 **(Figure 2A)**. These spheroids also displayed N-cadherin-positive membranes **(Figure 2B)**, indicative of the emergence of neural rosettes [6, 17]. In additionally, hCSs were generally positive for Pax6, reinforcing the fate specification of a cortical neural progenitor cell (NPC) population **(Figure 2B)**.

**Figure 2.**
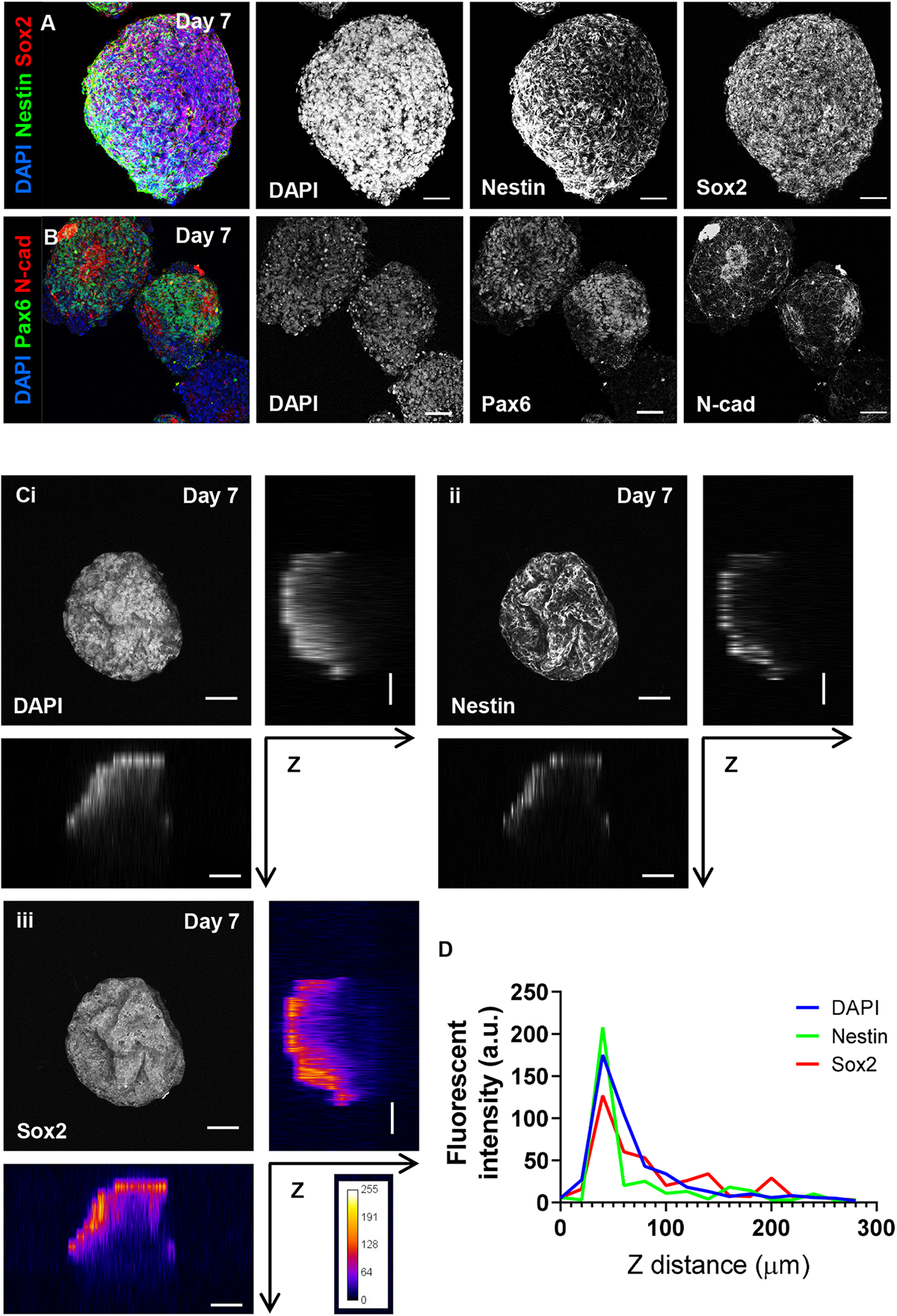
Confocal imaging of wholemount day 7 human cortical spheroids (hCSs) is limited at depth. **(A)** Representative confocal images of intact (wholemount) day 7 hCS showing strong immunostaining for the neuroepithelial/NPC markers Nestin and Sox2. **(B)** Representative confocal image of wholemount day 7 hCS demonstrates hallmarks of the emergence of neural rosettes, as determined by N-cadherin (N-cad) staining of apical membrane and surrounding Pax6 positive cells. **(C)** Maximally projected and orthogonal views of Z stacks of intact Day 7 hCSs acquired by confocal imaging. Confocal imaging displays a drop-off in fluorescence intensity correlated with acquisition depth (Z planes). **i)** DAPI, a non-antibody stain for DNA shows incomplete detection throughout the tissue-like structure. Similarly, **ii)** Nestin-positive and **iii)** Sox2-positive immunostaining displays even greater drop off in depth-detection, shown in **iii)** as a fluorescence-intensity heatmap. **(D)** Graphic depicting changes in fluorescent intensity (A.U.) with increasing imaging depth (μm) of **Ci-iii)**. Scale bars = 50 μm.

Although confocal microscopy was easily able to acquire z-stack images of superficial regions of hCSs, it was insufficient to acquire images of the entire spheroids. On average, we were only able to image up to 50-100 μm deep into hCS before significant signal drop-off occurred. This is demonstrated in **Figure 2C** where we have imaged a day 7 hCS of approximately 200 μm in diameter, stained for DAPI, Nestin and Sox2. Images were acquired as a z-stack – 15 images were acquired every 20 μm (1z step =0.4 μm using a 63x NA1.4 oil immersion objective) resulting in a total depth of 300 μm (**Supplemental Figure 2A-C)**. In all 3 channels, fluorescent signal towards the centre of the hCS tissue weakened dramatically at greater depth (>100 μm), as demonstrated by intensity profiles for each channel through the z-plane **(Figure 2C and D; Supplemental Figure 2D)**. This indicated either a poor acquisition of fluorescent signal via confocal acquisition or a failure of antibodies to fully permeate the tissue during preparation. However, fluorescent signal was similarly weakened beyond 100 μm in the DAPI channel, which is a highly penetrant small molecule fluorescent stain for DNA **(Figure 2C and D; Supplemental Figure 2D)**. This indicated that confocal microscopy was unable to image internal structures within intact spheroids.

We next characterised day 17 hCS, which were considerably larger than day 7 hCS. As day 17 hCSs ranged between 800-1200 μm in diameter (**Supplemental Figure 1C**), it was not possible to image intact day 17 spheroids using confocal microscopy. Consequently, day 17 hCSs were prepared as 20 or 60 μm cyro-sections before immunostaining and subsequent imaging with by confocal microscopy. Imaging of thin (20 μm) day 17 hCS cryo-sections demonstrated the expected neural rosette organisation of axial polarity with a lumen delineated by adjoining neural progenitor cells (**Figure 3A-D**). The inner lumen of neural rosettes was highlighted by the presence of tight junction proteins ZO-1 and N-cadherin (N-cad), as well as PKCλ-positive cells, indicating maintenance of adherent junctions by neural progenitors (**Figure 3A-C**). In the region immediately surrounding the inner lumen, nestin- and Pax6-positive cells were radially organised as expected for early NPCs (**Figure 3A & B**). DCX and MAP2-positive cells representative of the differentiation of NPCs into newborn and immature neuronal cells, could be seen constituting the outer layers of the neural rosette (**Figure 3C & D**). Overall, the description of neural rosette organisation and structure was similar to that previously described in 3D [2, 6, 17].

**Figure 3.**
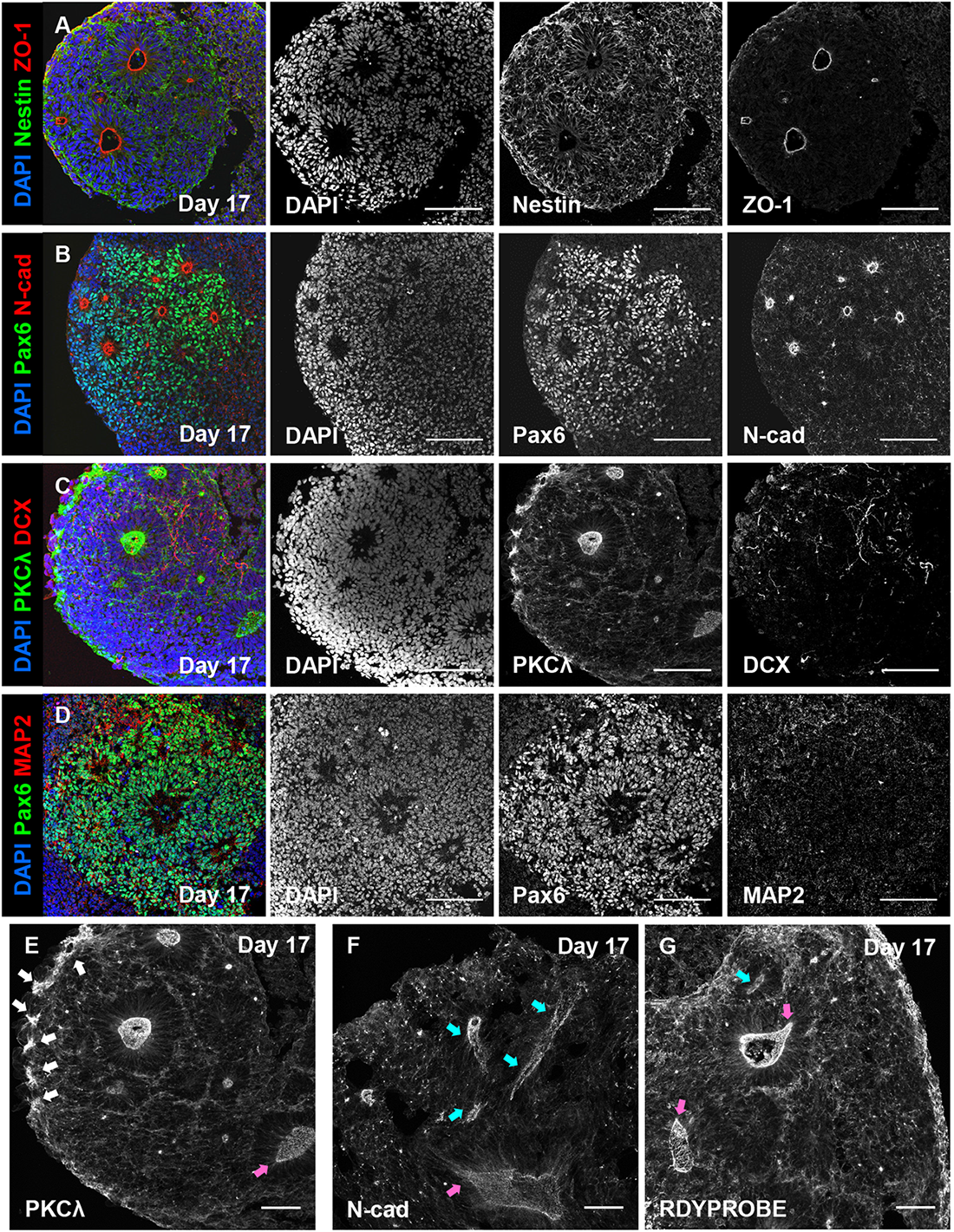
Confocal imaging of cryo-sectioned day 17 hCSs reveals incomplete acquisition of neural rosette 3D morphology. **(A-D)** Representative confocal images of sectioned (20 μm) day 17 hCSs; **(A)** ZO-1 staining of apical membrane is surrounded by apico-basal polarised Nestin filaments; **(B)** Pax6 positive cells self-organise around N-cad positive foci; **(C)** Immature DCX-positive neuronal processes are found outside of neural rosette structures; and **(D)** Map2-positive neuronal processes are also found to surround Pax6-positive NPCs organised into rosettes. **(E)** Confocal imaging of 60 μm sectioned day 17 hCSs express signs of cortical rosette formation. Newly formed PKCλ-positive membranes are seen at the edge of the tissue as indicated by white arrows. **(F)** Within thicker sections (60 μm) internalised tubular structures positive for N-cad are observed, suggesting that the inner lumen of neural rosettes may form into tube-like structures. **(G)** Phalloidin 488-positive actin filaments (RDYPROBE) show a concentrated organisation around ventricle-like structures. Magenta arrows in **E-G)** images highlight the ovoid formation of cortical rosettes with a pinched end, whereas blue arrows highlight formation of a tubular-like structure. Scale bars = 100 μm (A-D); 50 μm (E-G).

Interestingly, imaging of thicker sections (60 μm) occasionally revealed further insight into the emergence of rosette structures as well as greater detail on the potential morphology of the rosette lumen. For example, PKCλ, a putative organisational marker of neural rosettes, could at times be seen in pinched groups of cells at the outer layer of tissue (**Figure 3E white arrows**). These PKCλ-positive cells are thought to represent cells that internalise to subsequently form the inner lumen of rosettes [17, 29] In addition, when we imaged thick (60 μm) sections stained for N-cadherin, we were also able to occasionally observe rosette lumens as uneven, elongated or tubular structures (**Figure 3F**). This raised the possibility that the inner lumen of neural rosettes develops a tube-like morphology when spatially un-restricted in 3D and that we had serendipitously sectioned laterally through the inner lumen of a rosette **(Figure 3F; cyan arrow)**. A similar morphology was also observed in hCS sections stained for F-actin using an AlexaFluor 488-conjugated phalloidin stain (ActinGreen™ 488 ReadyProbes™ – referred to as RDYPROBE). RDYPROBE staining of f-actin, a major cytoskeletal component, was concentrated around the inner lumen of neural rosettes **(Figure 3G; cyan arrow)** similar to previous reports [17, 30]. However, f-actin staining could also be seen as polarised tube-like structures (**Figure 3G; magenta arrows**) similar to that seen with N-cadherin staining. Taken together, these data indicate that rosette lumens may develop into tube-like structures when grown in 3D, and that cryo-sectioning followed by confocal imaging is not sufficient to fully capture the morphology of these structures in sectioned spheroids.

### ALSM imaging of intact hCS

Having established that confocal microscopy was limited to imaging ~100 μm deep into intact spheroids, we next tested the ability of ALSM to image at greater depths. To this end, we generated day 30 hCSs, and stained them with RDYPROBE to label f-actin, and thus the structure of neural rosettes. An ALSM image of a day 30 hCSs (diameter = ~1.2 mm) was acquired as a single z-stack. A raw output max intensity Z-projection (z-stack = 500 images – 600 x 600 x 600 μm) can been seen in **Supplemental Figure 3Ai (Raw image)**. This image stack subsequently underwent deconvolution using a Richardson-Lucy algorithm [31, 32] specific to the ALSM (MSquared Cubes software). Following this image processing, multiple neural rosettes with tube-like structures could be observed within the hCS **(Supplemental Figure 3Aii – Deconvolved Image)**. However, we noted that some non-specific staining on the surface appeared very bright leading to features of interest appearing dimmer. Therefore, we removed the images of the outer layer of tissues from the 3D image stack. This removed highly stained cells at the edge which had a higher fluorescence than internal structures, and thus allowed for a clearer visualisation of internal rosettes **(Supplemental Figure 3B)**. Multiple neural rosette lumens with different shapes and sizes could be observed in both 3D renders and max projection images (**Figure 4A & B**). Larger rounder rosette lumens were observed towards the middle of the tissue whereas smaller extended tube-like rosettes can be found towards the edge of the tissue (**Figure 4A & B**). These structures had diameters ranging between 20-50 μm, and lengths between 20-200 μm.

**Figure 4.**
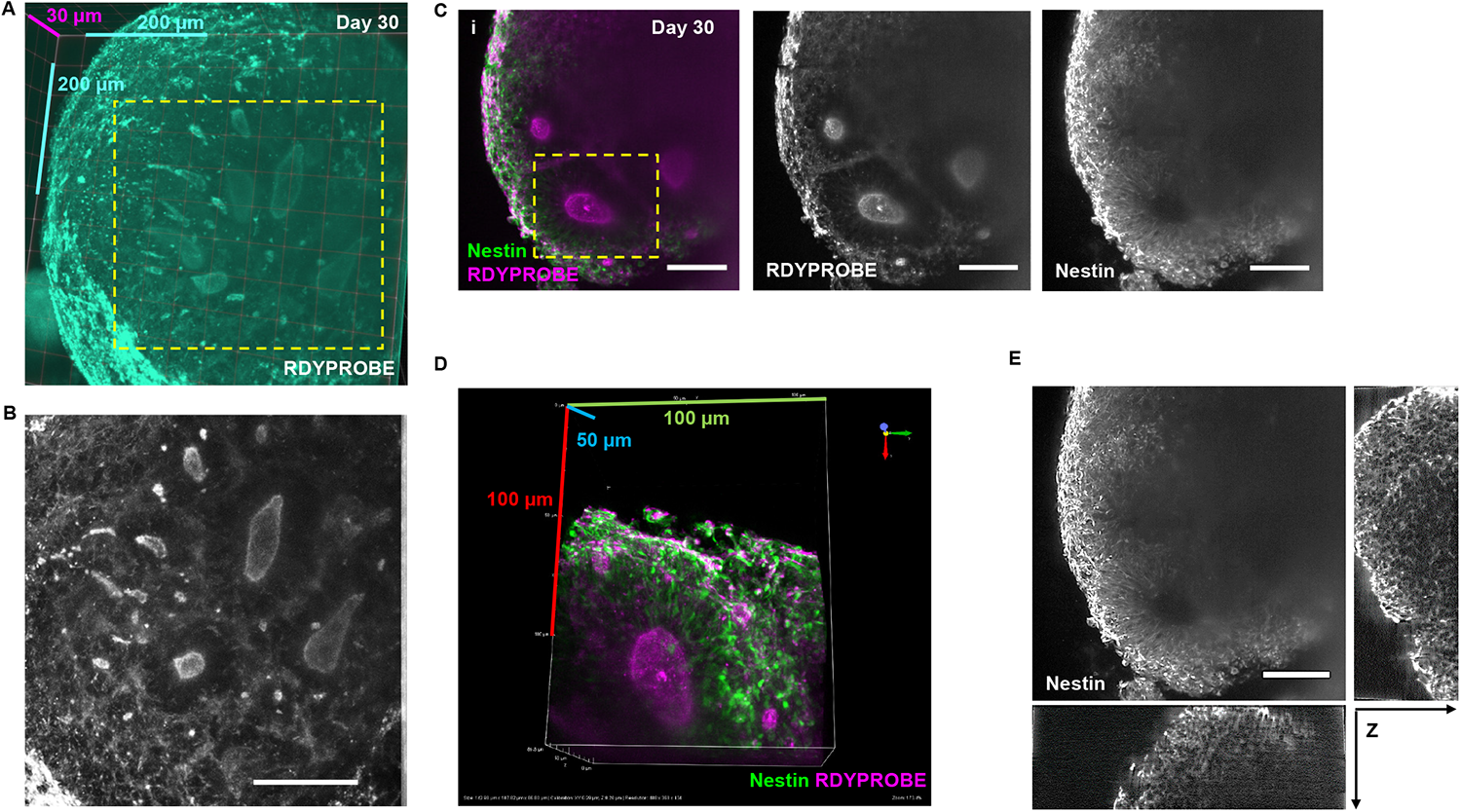
Visualising internal structures of hCSs using Airy-beam light sheet fluorescent microscopy. **(A)** Clearvolume 3D representation of multiple internalised rosette lumen of differing sizes shown by RDYPROBE staining (F-actin) with dimensional scale. **(B)** Magnified region (A, dotted line) showing characteristic ovoid or tubular shaped lumens expected of a ventricle-like structure. **(C)** Composite **i)** and individual channel max-projected images of **ii)** RDYPROBE (F-actin) staining and **iii)** Nestin-positive filaments around apical membranes, showing the presence of multiple large rosettes, Scale bars = 100μm. **D)** 3D render of magnified region from **Ci** (dotted line) displaying of a singular rosette lumen co-stained for radial Nestin filaments and RDYPROBE (F-actin) with dimensional scale. **E)** Orthogonal views of stained tissues permit examination of a cortical rosette in all axes. Scale bar = 50μm.

We next imaged a day 30 hCS that was stained with RDYPROBE and immunostained with the NPC marker Nestin to assess signal intensity deep within day 30 hCSs **(Figure 4C and Supplemental Figure 3C**). As expected, F-actin staining revealed several neural rosette lumens within the spheroid. High magnification zoom-ins of f-actin enriched apical membrane showed that Nestin-positive cells were organised radially around the central lumen **(Figure 4C)** with Nestin filaments being polarised and projecting away from the central lumen **(Figure 4C)**. However, examination of individual channels revealed differences in signal intensity and image clarity at depth **(Figure 4C-E)**. F-actin staining could be seen though most of the spheroid **(Figure 4C & D; Supplemental Figure 3D)**, with signal intensity dropping when reaching the inner most regions of the spheroid. Simiarly, the signal intensity from Nestin staining substantially dropped at more superficial depths compared to the f-actin stain, with a loss of resolution at depth **(Figure 4D & E)**. Thus, the organisation of Nestin-positive cells and f-actin apical lumens rosette lumen was limited at depth (>200 μm). Despite this, orthogonal examination displayed ALSM maintains a much greater resolution and clarity compared to confocal when imaging at depth **(Figure 4E)**. Together these data demonstrate that the ALSM can rapidly (500 z slices within ~20 sec) acquire multi-channel 3D volumes, but with limited resolution when imaging hCS at depth.

### Optimised immunohistochemistry and tissue clearing improves ALSM imaging of deep structures within hCS

Studies have shown that effective imaging at depth can be enhanced by clearing the 3D tissue and optimising immunostaining protocols before LSM imaging [19, 33, 34]. As our day 30 spheroids were approximately 2mm in diameter (**Supplemental Figure 1E**), we reasoned that optimizing our immunostaining protocol would improve antibody penetration in 3D tissue as previously described [25]. As, it is well understood that lipids cause light scattering which interfere with deep tissue imaging, we further reasoned that cleared our hSCs so that the lipids were replaced with a clear water-soluble matrix, would reduce light scattering and therefore improve fluorescent signal within spheroids [24, 25]. Optimised immunostaining was achieved by using glycine and DMSO at low concentrations which improve pH and antibody stability for longer incubation periods, while using Tween-20 to help make cells more porous for better antibody penetration [25]. Next, hCSs underwent tissue clearing using the ScaleS4 method, a rapid aqueous-based tissue clearing approach that which takes 4 hours [24].

Cleared hCSs were immunostained with Sox2 (NPC marker) and ZO-1 as a marker of the apical membrane of rosettes (**Supplemental Figure 4A)**. Unlike non-cleared hCSs, antibody signal could be detected throughout the 600 x 600 x 600 μm captured image without any obvious loss of fluorescent signal (**Figure 5A-C; Supplemental Figure 4B and Supplemental Movie 1)**. Both ZO-1 positive apical membranes as well as Sox2-positive cells could be easily detected in optical sections at various levels within the 500 image z-stack **(Supplemental Figure 4B)**. In 3D renders of the imaged volume **(Figure 5B)**, both ZO-1 and Sox2 could be seen at different depth within the spheroid **(Figure 5C & Supplemental Video 1)**.

**Figure 5.**
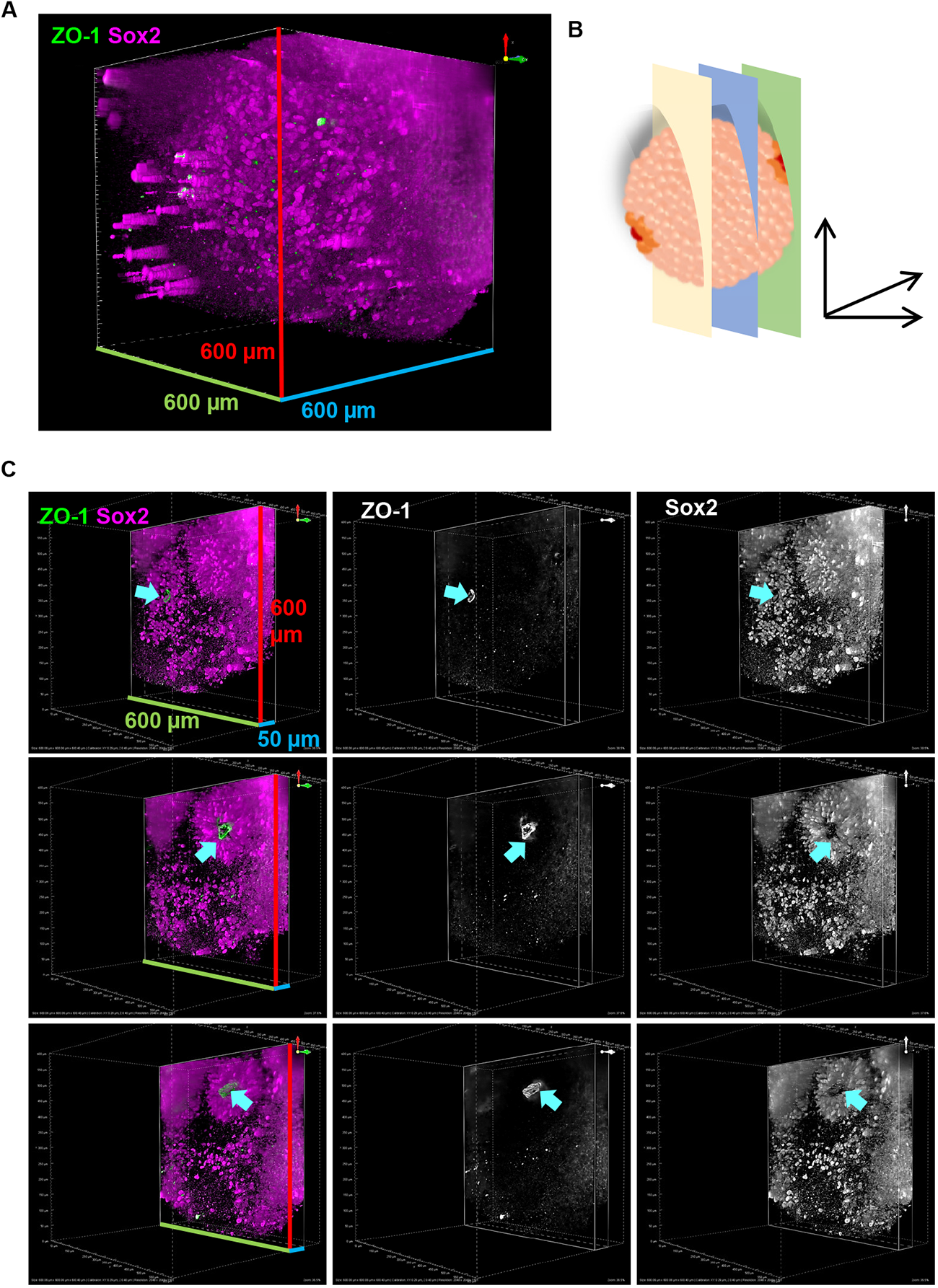
ALSM imaging of cleared hCSs. **(A)** Representative 3D volume image (600 x 600 x 600 μm) of cleared day 30 hCS immunostained with ZO-1 (green) and Sox2 (magenta) and imaged by ALSM. **(B)** Schematic image of ALSM optical sections of cleared hCS shown in (**C**). **(C)** ALSM optical sections (600 x 600 x 50 μm) at 450, 500 and 550 μm depth, of cleared hCS shown in (**A**). Cyan arrows indicate ZO-1-positive rosette lumen with surrounding radially organised SOX1-positive NPCs. Green and red bars indicate scale in X and Y planes (600 μm) – blue scale bar indicates scale in Z plane (600 μm in (**A**) and 50 μm in (**C**)).

Consistent with expected results, Sox2-positive cells were organised in a radial manner around ZO-1 positive rosette lumens (**Figure 5C; Supplemental Figure 4B)**. Together these data indicate that the combination of an optimised immunostaining protocol with an aqueous clearing method vastly improves imaging depth in hCS.

### ALSM imaging of hCS reveals 3D morphology of neural rosette inner lumen

Confocal imaging of sectioned day 17 hCS indicated that the inner lumen of neural rosettes may develop large spherical or tubular lumens (**Figure 3C-E**). However, it was not possible to gain an understanding of the 3D morphology the central lumen by confocal microscopy. Conversely, ALSM imaging of day 30 hCS stained with ZO-1 (**Figure 5**) indicated that it may be possible to visualise the intact inner lumen of a rosette. To this end, we generated 3D renders of the ZO-1 channel from ALSM imaged day 30 cleared hCSs. Multiple ZO-1 positive structures could be detected within a single hCS (**Figure 6A**). Morphological measurements of ZO-1-positive rosette lumens revealed that they varied in size and volume (**Figure 6A & B**). Most ZO-1-positive lumens (>95%) had volumes ranging between ~150 to < 10,000 μm^3^: 75% of measured lumens were less than 1000 μm^3^ **(Figure 6C)**, and a surface area that ranged between ~200 to > 10,000 μm^2^; again the majority of lumens (~66%) were less than 1000 μm^2^ **(Figure 6C)**. A single large rosette lumen could be readily observed **(Figure 6E & F; Supplemental Figure 5 A & B)**. This structure had volume of 192,484.21 μm^3^ and corresponding surface area of 146697.61 μm^2^. It was also possible to extract additional morphological parameters including length at the longest axis and shape factor, a measure of sphericity (**Supplemental Figure 5 C & D)**. Finally, by focusing on a single large rosette lumen, it was possible to gain an appreciation of the radial organisation of SOX2-positive NPCs surrounding the inner lumen of rosettes in 3D **(Figure 6G; Supplemental Figure 5 E and Supplemental Movie 2)**. Taken together, these data demonstrate that following an optimised immunohistochemical and tissue clearing approach, that it is possible to gain insights into the 3D structure and organisation of neural rosettes and surrounding NPCs in hCSs.

**Figure 6.**
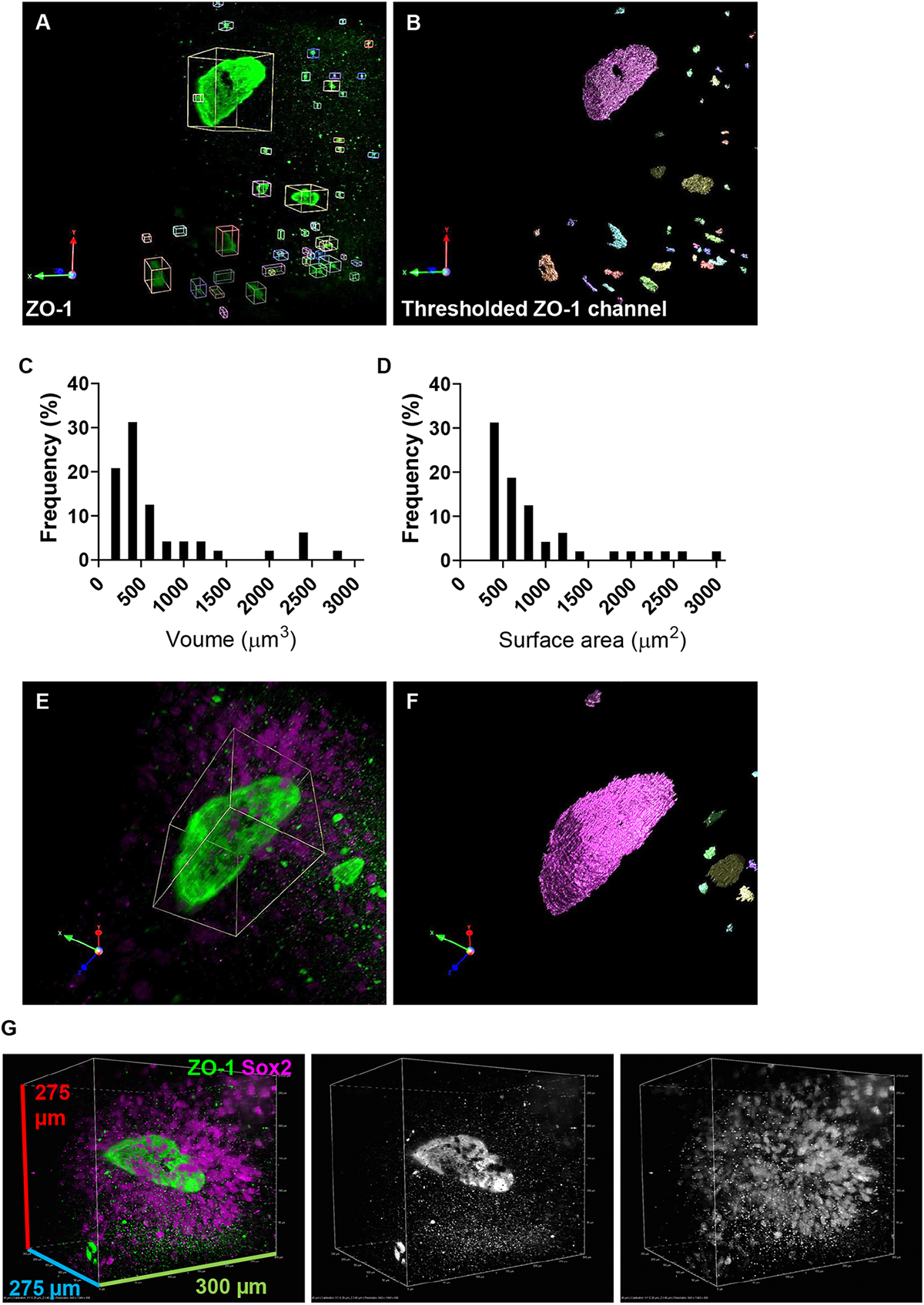
Visualisation and quantification of neural rosette lumen morphology in intact cleared hCSs. **(A)** Representative 3D volume images of ZO-1-posiitve apical lumen membranes of cleared day 30 hCS. **(B)** Rosette lumens were defined as objects in Volocity (Perkin Elmer) by thresholding volumetric regions based on ZO-1 channel. Objects with volumes of less than 150 μm^3^ were discarded. **(C & D)** Frequency distribution of neural rosette lumen volume (μm^3^; **C**) and surface area (μm^2^; **D**). Measurements of these morphological parameters revealed a single neural rosette with a lumen with a volume of 192,484.21 μm^3^ and surface area of 146697.61 μm^2^. **(E)** Digital zoom of largest neural rosette lumen. Green channel indicates ZO-1-psotive apical membrane; magenta channel indicates Sox2-posiive NPCs surrounding central lumen. **(F)** Threshold image of ZO-1-positve lumen demonstrating morphology of structure in 3D. **(G)** Representative 3D render of largest neural rosette lumen demonstrating the radial organisation of Sox2-positve NPCs surrounding ZO-1-positive (green) rosette lumen.

## Discussion

The ability to generate 3D brain organoids from somatic cell-derived human hiPSC is a great tool to further our understanding of human brain development in health and disease [2–5, 7, 10, 11, 18]. However, many conventional tissue processing and imaging modalities are not optimized for the preservation and imaging of intact organoids. This result in the loss of the significance and understanding of the cellular processes, organisation and complexity within a three-dimensional environment [19]. Here we have optimized and immunohistochemical and tissue clearing approach in combination with ALSM imaging that enables for the rapid and high resolution imaging of intact hCS. Using this approach we were able to visualise and quantify the three dimensional cellular structure of the inner lumen of neural rosettes. This structure is described as an in vitro correlate of the human neural tube which begins to form at gestational week 4 and is the source of the entire central nervous system [17]. As the formation or maintenance of structure of neural rosettes have been reported to be altered in autism and related neurodevelopmental disorders [8, 9, 35], being able to examine the cellular organisation and morphology of neural rosettes in 3D may offer insight into pathological mechanisms that contribute to the likelihood of developing such disorders.

Conventional methods to study brain organoids such as hCS are often hampered by a number of limitations such as the inability to acquire images of cellular structures and organisations within intact specimens. For example, we found that even imaging of intact hCS with a diameter of ~300 μm resulted in a rapid loss of fluorescence signal approximately 100 μm into the tissue. This was not due to an issue of antibody penetration as the observed signal loss was also observed when imaging DAPI, a non-antibody stain. We were able to overcome this limitation by using an ALSM imaging modality. Advantages of using this form of LSM include the ability to rapidly acquire volumetric images, a large FOV [21]. Moreover, due to the nature of the Airy-beam, this form of LSM can produce high-resolution images with less photo damage to the specimen [34]. Imaging of hCSs significantly improved acquisition of fluorescent signals at depth – multi-channel volume imaging of 600 x 600 x 600 were acquired within 20 seconds. This allowed visualization of the inner lumen of neural rosettes in 3D in manner not possible using confocal microscopy. Using the f-actin stain RDYPROBE, we observed the ovoid or tubular shaped lumens of neural rosettes. It was also possible to observed radial arrangement of NPCs cells around the lumens. This was our first evidence of the 3D shape of neural rosettes and surrounding NPCs.

Despite this improved ability to image neural rosettes within intact hCSs, we still observed a loss of signal was still evident at depth, albeit much less that than observed following confocal imaging. It is well recognised that the presence of lipids within tissue samples cause a scattering of light, and therefore impedes effective volume imaging [19, 22, 24, 33]. In addition, uneven, or incomplete penetrance of antibodies could also contribute to this issue [25]. Our approach to address the issue of uneven or imcmplete antibody penetration was to use the optimized immunostaining approach described in the FACT clearing protocol [25]. However, in this protocol, tissue clearing takes 1 week followed by 4 days for antibody staining. In order to shorten this duration, we used the optimised immunostaining protocol in FACT (4 days) in conjunction with a more efficient and quick clearing method, ScaleS4 which takes 4 hours [24]. By combining this approach with ScaleS4 clearing, our tissue preparation time was reduced to less than half (~5 days) of the FACT protocol. This is also much less compared to other established protocols such as CLARITY (~20 days) and CUBIC (~11 days) [22, 24, 25]. As FACT and ScaleS4 were both aqueous-based protocols, we found they were quite compatible when used together. This optimized tissue clearing approach further improved ALSM imaging of intact hCS. We visualised the inner lumen, as well as neuroepithelial cells surrounding the lumen to provide a representation of the 3D structure of the neural rosette. To our knowledge, this is one of the first studies to examine the lumen of neural rosettes in 3D. From the cleared images of neural rosettes, we were able to clearly delineate the edges of the lumen and create a 3D reconstruction of the shape of the lumen. This allowed us to quantify the size of the lumens and found that most lumens in a given spheroid had volumes less than 1000 μm^3^, while only few had volumes larger than 10,000 μm^3^.

The presence of a single larger lumens corresponded is similar to that previously reported [17]. The formation of a single large rosette is considered an essential feature of normal brain development and critical for establishing typical tissue cytoarchitecture and normal differentiation of neuroepithelial cells during brain development [17]. Consistent with this, it was possible to observe the radial organisation of NPCs around the large rosette in 3D[2, 5–7]. Thus, this approach suggests that it is possible to examine typical cellular maturation observed in neurodevelopment in 3D. Thus, this combined approach would be a good cellular model to study neurodevelopmental conditions such as autism.

## Limitations

The ALSM system used in this study has a working distance of 3.5 mm due to the physical room between the illumination and detection objectives. Furthermore, while a day 7 hCS could fit fully into the FOV, only a part of the day 30 tissues (600 x 600 x 600 μm) could be imaged. A number of alternative forms of LSMs have been developed that could be used as an alternative [19, 36]. Nevertheless, despite these limits, a rapid acquisition time (~20 seconds for 500 z-slices) allows for quick repositioning of tissues to new fields of view or different ranges of depth. Reduced excitation during acquisition allows extended periods of imaging, as photo-bleaching is minimal, making the system suitable for live-imaging of developmental processes, as it does not require clarification of tissues nor sectioning.

In this study we have generated hCS brain organoids as an in vitro model of cortical development. However, this protocol is variable, often producing organoids of various sizes, and moreover, only generates cells representative of the dorsal forebrain. A number of protocols have been described for the generation of 3D brain organoids with less variability and of different brain regions in addition to the generation of ‘fused’ organoids that model multiple brain regions [1–3, 14–16]. Such approaches could be used to reduce variability and would facilitate the modelling and examination of cellular organisation, process, and interaction between multiple brain regions.

The key problem being faced by the field of volumetric acquisition (including the use of ScaleS or CLARITY) is the need for standardised methods of volumetric quantification, in order to fully extrapolate the vastly powerful data obtained [19, 22, 26, 36]. Due to current popularity of 2D cell culture and sectioned tissues, many means of quantification disregard the possibility of a third spatial dimension. However, a number of programs, including open-source programs, that include high-through acquisition and automated quantification of 3D structures are being developed [36–38].

## Conclusion

In summary, the combined immunohistochemical, tissue clearing and ALSM imaging approached described in this study represents a significant step forward from conventional confocal microscopy. Using this approach, it is possible to visualise cellular structures critical to human brain development in the hCS in virto model in 3D. From imaging 3D spheroids, we have observed the ovoid, tubular structure of the lumens, and the radial arrangement of differentiating neural progenitor cells within the neural rosette. We also observed larger neural rosettes were fewer in number in a spheroid, approaching the singularity that is observed in vivo during neural tube formation. This was evidence that this method could be used to study typical human brain development, and characteristics of neural rosette formation could be used to study brain development in neurodevelopmental conditions such as autism.

## Methods

### Study aim and design

The aims of this study encompassed two core objectives: 1) comparison of conventional tissue processing and image acquisition methods against a novel methodology of presenting intact 3D tissue for volumetric acquisition using an ALSM; and 2) to determine whether clarification of tissue would improve acquisition of neural rosettes deep within hCSs. To this end, hCSs were differentiated from 3 previously well characterise hiPSCs [shum et al., 2020] – CTR_M1; CTR_M2 and CTR_M3 – up to 2 clones per line were utilized in the study. For all experiments, hCSs were generated from each line (1-2 clones); images presented in figures are representative of structures observed from all lines. Between 2-5 independent batches of hCS were generated for each experiment where a batch was defined as an independent differentiation from hiPSCs.

### HiPSC maintenance

HiPSCs from 3 typically developing individuals (CTR_M1; CTR_M2 and CTR_M3) were generated as previously described as part of EU-AIMS and STEMBANCC EU IMIs [cocks, deans, shum et al]. Ethical approval for the generation of hiPSCs were approved under the PiNDS study (REC: 13/LO/1218). HiPSCs were grown and maintained in E8 medium (Life Technologies) with E8 supplement (Life Technologies). HiPSCs were passaged by first dissociating attached cells from the plate, by incubating them in Versene (Thermo Fisher 15040066) for 4-5 minutes in 37°C incubator. Versene helps gently lift edges of iPSC colonies, after which it was replaced with E8 and cells were lifted using a cell lifter. Free-floating cells were re-plated onto fresh geltrex-coated (Thermo Fisher 12760013) plates. For maintenance, cells were re-plated at ~70% confluency. For starting a new experiment, cells were re-plated at ~100% confluency.

### Generation of human cortical spheroids

HiPSCs were differentiated into human cortical spheroids (hCS) using methods published by Pasca *et al*. [2]. In this method dual SMAD inhibition was performed using KOSR media supplemented with 10 μM ROCKi and SMAD inhibitors: 5 μM dorsomorphin (Tocris) and 10 μM SB431542 (Tocris) to direct hiPSCs towards a cortical lineage. Cultures were maintained in 5% CO2 at 37°C for 48-hours period to promote formation of spheroids. From day 2 until day 4, media changes were performed daily with fresh KOSR media supplemented with 5 μM Dorsomorphin and 10 μM SB431542. From day 5 until day 25 (or when hCSs were harvested), culture media used was B27 minus Vitamin A (Life Technologies) media plus 20 ng/mL of recombinant human EGF and recombinant human bFGF (both Preprotech). From day 25, B27 minus Vitamin A media with 20 ng/mL of recombinant human BDNF (Preprotech) and recombinant human NT3 (LifeTechnologies) was used. Media was changed every 24 hours during the first 15 days and once every 48 hours thereafter. Spheroids were harvested at days 7, 17 or 30.

### Sample preparation for confocal microscopy

Cortical Spheroids harvested at **Day 7** were fixed in 4% formaldehyde in 4% sucrose/PBS for 45 minutes then washed three times with PBS. Once fixed, spheroids were permeabilised in 0.3% Triton X100/PBS for 30 minutes, before permeabilisation-blocking for an hour in 2% NGS in 0.3% Triton X100/PBS. Subsequently spheroids were incubated in a solution containing 1° antibodies, 2% NGS and 0.1% Triton X100/PBS at 4°C for a 48-hour period. Spheroids were then washed three times in PBS before being incubated overnight in a solution of 2° antibodies, 2% NGS and 0.1% Triton X100/PBS in a lightproof container at 4°C. Post-incubation spheroids were washed twice in PBS before DAPI stain was applied. Prepared samples were mounted intact onto Superfrost plus slides (VWR, 631-0108P) by suspending in Mowiol mounting media (Sigma Aldrich, 81381) and surrounding with silicon grease (to protect tissue from high pressures) before applying glass coverslips (VWR, 631-1574) and leaving for 48 hours to set (Figure 6b). Prior to imaging using confocal microscopy, slides were stored at 4°C in the dark. Spheroids harvested at **Day 17** were fixed in 4% formaldehyde/PBS for 1 hour, followed by three PBS washes before incubation in 30% sucrose solution (weight/volume) overnight. Once spheroids had normalised they were placed into moulds and cryopreserved in OCT or M1 freezing media via snap-freezing on dry ice, before storage at −80°C. Cryopreserved tissues were cryosectioned using a Leica CM1850 cryostat at 20μm, 40μm and 60μm section depth prior to mounting on Superfrost plus slides for fluorescent immunohistochemistry. Sample slides were treated via the same process as post-fix 2D cell culture coverslips, before being mounted with Mowiol and a glass coverslip. Prepared slides were stored in darkness at 4°C, before image acquisition via confocal microscopy. Compared to Day 7, Day 30 spheroids were significantly larger (Figure 5c and 5e) and thus required longer treatment to ensure penetrance of washes into deep tissue. **Day 30** spheroids were fixed in 4% formaldehyde/PBS for 2 hours, followed by rinsing with three PBS washes, before a permeabilisation-blocking step of 10% NGS in 0.3% Triton X100/PBS for 1 hour 30 minutes. Permeabilised spheroids were incubated for 48 hours at 4°C in a 1° solution, consisting of 1° antibodies in 10% NGS and 0.1% Triton X100/PBS. Subsequently spheroids were washed three times with PBS and incubated in a 2° solution consisting of 2° antibodies in 10% NGS and 0.1% Triton X100/PBS overnight at 4°C, protected from light. Stained spheroids were washed thrice in PBS before being transferred to 0.02% NaN3/PBS for long-term storage to prevent microbial growth and tissue degradation. Once ready for imaging, tissues were immersed in ~50°C 0.1% low melting point agarose and set onto specialised mounting slides for light sheet acquisition (Figure 6c). Tissues could be reclaimable by disintegrating the agarose-matrix bead in PBS with time, thus permitting reimaging.

Permeabilised samples were incubated in solution, consisting of primary antibodies **(Supplemental Table 1)**. Superfrost plus slides using Prolong Gold mounting media (Thermo Fisher, P36934) was used to reduce photo bleaching of fluorophores and left in darkness at RT overnight to set, before imaging via confocal microscopy.

### Sample preparation for ALSM imaging of intact hSC

#### Non-cleared spheroids

The cortical spheroids were first washed in PBS for 3 minutes and then fixed in 4% formaldehyde in 4% sucrose-PBS. The fixation time was 45 minutes for day 30 spheroids. Once the spheroids were fixed they were washed again in for 3 minutes (3 times) in PBS and stored in sucrose 30% (weight/volume) the sucrose sinking improves their preservation. After sucrose sinking, the spheroids were washed three times in PBS for three minutes each time prior to the permeabilisation step. In order to permeabilise the tissue the antibodies could reach their epitopes within the cell membrane and to prevent non-specific staining the samples were incubated for 60 minutes in permeabilisation-blocking solution of 2% normal goat serum (NGS) in 0.1% Triton X100 and PBS. Then the spheroids were incubated in a solution of permeabilisaton-blocking solution containing the primary antibodies for 48 hours. The samples were washed three times for three minutes in PBS and incubated with permeabilisation-blocking solution containing the secondary antibodies and HOECHST (nuclear staining) for two hours and kept in darkness to prevent bleaching of the fluorophores.

#### Cleared spheroids

The cortical spheroids were first washed in PBS for 3 minutes and then fixed in 4% formaldehyde solution in 4% sucrose-PBS overnight at 4°C. Fixation solution was then removed and stored in sucrose 30% (w/v) overnight at 4°C. Spheroids were then permeabilised and blocked using a perm/block solution (0.6M glycine, 0.2% Triton X-100, 2% goat serum, 20% DMSO in PBS) overnight at room temperature (RT). Samples were permeabilised for this length of time in order to facilitate antibody penetrance into large tissues. The samples were washed twice in PBST for 1hr each time at RT. Primary antibody was made using an ‘antibody solution’ (0.2% Tween-20, 5% DMSO, 2% goat serum, 0.01% sodium azide in PBS). Spheroids were incubated in antibody solution for 2 days. After 2 days, spheroids were again washed in PBST for 1hr each at RT. Secondary antibody solution with added Hoechst (for nuclear stain) was made using the same ‘antibody solution’ as before. This was incubated similarly for 2 days at RT, protecting from light. Following Secondary staining, cells were washed with PBST, and stored in PBS before clearing and mounting for ALSM imaging.

### Clearing and mounting of intact hCS

To mount samples a set of homemade sample mounts were created by bonding a 46mm square weighing boat (Fisherbrand Cat No. 12942860) to a standard 75mm microscopy slide using cyanoacrylate glue (**Supplemental Figure 6**). A silicone rubber plinth was constructed by first casting silicone rubber (9 parts reagent to 1 part catalyst) in a flat vessel (petri dish) with a textured base (60 grit sandpaper) to a depth of approximately 5mm and left to cure for at least 24 hours. Once this had set a 4mm cylinder was punched using a hollow hole punch tool. The cylinder was bonded to the middle of the weighing boat with the textured surface facing upwards using cyanoacrylate glue and left to set for a few hours before use. The silicone rubber cylinder formed the imaging plinth for samples to be positioned on allowing lenses to lower around plinth and sample without collision to base of the weighing boat (**Supplemental Figure 6**).

In order to place whole hCSs on to the imaging plinth 1.2% w/v agarose in water was melted and a small volume (less than 50 μl) was placed on to the plinth and the spheroid was positioned in the top of the still liquid agarose shortly after allowing agarose to solidify around the spheroid and holding it still for imaging.

Mounted spheroids were then cleared using the ScaleSQ5 protocol published by Hamma et al. 2015[24]. In this method, clearing was performed by incubating hCSs into ScaleSQ(5) solution (**Supplemental Table 2**) for 2 hours at 37°C. this was followed by submersion of the cleared tissue in mounting media ScaleS4(0) (**Supplemental Table 2**) for at least 2 hours at room temperature. The mounting media had a refractive index (RI) = 1.437.

### Confocal microscopy

A Leica laser scanning confocal microscope (TCS-SP5) was used to image Day 7 and 17 2D cultures, as well as Day 7 whole-mount and Day 17 cryosectioned spheroids. Leica’s LAS AF software was used to set up and acquire multiple channels of singular planes and Z stacks at a resolution of 1024×1024px. 405nm (DAPI; blue), 488nm (green), 561nm (red) and 633nm (far-red) lasers were used to generate images in colour channels. Smart gain was used to adjust the different channels to a similar intensity to highlight fluorescent staining. 2D cultures were acquired at 63x magnification in oil immersion (NA 1.4), whereas 3D cultures were imaged at 40x magnification in oil immersion (NA 1.3 to check) to encompass a wider field of view. A line average of three was used in each channel during point scanning to reduce appearances of random background signal caused by light scattering, therefore increasing the quality of acquisition.

### Lightsheet microscopy

An upright Aurora™ M Squared single photon Airy beam^1^ light-sheet microscope was used to perform the light-sheet imaging on the organoid samples. The fluorophore excitation was delivered with a 10x water immersion illumination objective (Olympus UMPLNFLN 10x NA 0.30) and emission was captured orthogonal to the excitation plane using a 20x water immersion detection objective (Olympus UMPLNFN 20x_NA 0.50) to provide a lateral resolution of 670 nm. The emission light passed through a filter wheel before images were captured using a sCMOS camera (Orca Flash 4.0 v2) with a FOV of 600 μm. Three-dimensional volumes were acquired in step sizes of 0.4 μm by XZ translation of the sample through the focal plane and outputted stacks were deconvoluted to provide an axial resolution of ~870 nm. Samples were excited using a 405 nm (DAPI; blue), 488 nm (green), 561 nm (red) and 640 nm (far-red) lasers and camera exposure rate adjusted to generate optimal S/N for multi-colour images. Objective working distance was 3.5mm which allowed ample space for positioning of samples.

### Image acquisition with Lightsheet microscopy

#### Preparation and collection of PSF for deconvolution / image processing with tetraspeck beads

In order to process the data with the M-Squared Lasers Deconvolution software a volume of data containing fluorescent microspheres was collected with identical optical and spatial parameters (wavelengths for excitation and emission as well as xy pixel size and z step size) as the collected specimen data to be able to produce a calibrated point-spread function (PSF). To produce a suspension of fluorescent microspheres adequate for this purpose 1.2μl of stock solution of 0.5μm fluorescent microspheres (Invitrogen Tetraspeck 0.5μm – Cat No. T7281) was added to 100μl of molten agarose (1.2% w/v) and the mixture was completely mixed by pipette action before dispensing. 50μl of the mixture was drawn and a dome of agarose was placed on a clean sample mount and allowed to solidify before continuing. Mounted bead suspensions were then placed in the corresponding imaging medium (PBS or ScaleS0(4)) and either used immediately (PBS) or left overnight. For samples or beads in ScaleS0(4) at least one further solution change was then made a few hours before imaging to allow solutions to properly equilibrate and allow any residual water from the agarose to be completely removed.

Acquisition of bead fluorescence volumes was performed by positioning the bead preparation in the focal plane of the lenses with the motorised stage and first checking excitation beam was visible and focussed by halting scanning mirrors to produce a static beam image and observing position and shape of formed image and adjusting focus of excitation and detection objectives so that the correct focussed beam shape was observed. Once this was satisfied the scanning mirrors were reactivated and images were collected with excitation power and camera integration time sufficient that the peak image intensity of beads were measured to be at least 10% of the total camera bit depth (16bit – 65535) producing sufficient signal to noise ratio for software to automatically locate beads from the data set. This criterion was met for each combination of excitation wavelength and emission filter combination required to image a sample and a volume of at least 100μm depth of images spaced 0.4μm apart in focus. The stage was translated in the yz axis with a dedicated stage driver for this axis as this was the perpendicular direction to the focus of the detection objective. This data was analysed and processed by software to obtain the PSF of the imaging setup to be used for deconvolution. This generated a unique PSF and calibration file used on each date of data acquisition and used to deconvolve datasets of sample fluorescence.

#### Sample Image Acquisition, Deconvolution and Presentation

Samples once mounted were positioned in the focus of the objectives and data were collected by translation of the motorised stage using the yz driver. Laser power and camera integration times were set to provide sufficient contrast and intensity aiming to minimise total integration time while not compromising signal. The raw data were then passed to M-Squared Lasers proprietary deconvolution software to process for deconvolution and removal of airy pattern in data. Richardson-Lucy deconvolution was processed with at least 100 iterations and a final datafile is produced for each channel. These were then combined in Nikon Elements (version 5.02) to overlay channels and produce 3 dimensional renders as shown in figures. In some instances, Volocity 6.3.1 (Perkin Elmer) or ClearVolume (ImageJ plugin – https://imagej.net/ClearVolume) were used to produce 3D renders which was chosen only for convenience at the time.

Morphological measurements of lumen volume (Figure 6) were performed using Volocity 6.3.1 (Perkin Elmer) where an object is defined by use of the built-in “find-objects” method and selecting an intensity value to set the threshold for volumetric regions based on the channel used. An arbitrary volume of more than 150 μm^3^ were chosen from the dataset and objects smaller than this were discarded.

## List of abbreviations

ALSM: Airy light-sheet microscope
FOV: Field of view
FWHM: Full width at half maximum
hCSs: Human Cortical Spheroids
hiPSC: Human induced pluripotent stem cell
LSM: Light sheet microscopy
N-cad: N-cadherin
NPC: Neural progenitor cell

## Declarations

### Ethics approval and consent to participate

Ethical approval for the generation of hiPSCs were approved under the PiNDS study (REC: 13/LO/1218).

### Availability of data and materials

Data sets are available upon reasonable requests.

## Competing interests

JS, NH and RF are all (or were) employees of M Squared Life Ltd.

## Funding

The study was supported by grants from the European Autism Interventions (EU-AIMS) and AIMS-2-TRIALS: the Innovative Medicines Initiative Joint Undertaking under grant agreement no. 115300, resources of which are composed of financial contribution from the European Union’s Seventh Framework Programme (FP7/2007-2013) and EFPIA companies’ in kind contribution (DPS, SB-C); StemBANCC: support from the Innovative Medicines Initiative joint undertaking under grant 115439-2, whose resources are composed of financial contribution from the European Union [FP7/2007-2013] and EFPIA companies’ in-kind contribution (DPS); MATRICS: the European Union’s Seventh Framework Programme (FP7-HEALTH-603016) (DPS). In addition, funds from the Wellcome Trust ISSF Grant (No. 097819) and the King’s Health Partners Research and Development Challenge Fund, a fund administered on behalf of King’s Health Partners by Guy’s and St Thomas’ Charity awarded to DPS; the Brain and Behavior Foundation (formally National Alliance for Research on Schizophrenia and Depression (NARSAD); Grant No. 25957), awarded to DPS, were used to support this study. ACV acknowledges funding support from the Royal Society [RG130610]

## Authors’ contributions

DA, GC, JC, EV-A, JS, NH all carried out experiments, data acquisition and analysis; MVY, ACV and DPS provided technical advice; RF provided unique resources; SB-C, ACV and DPS provided funding; DA, GC, JC, NH ACV and DPS wrote the manuscript; DPS oversaw the project.

## Acknowledgements

We thank the Wohl Cellular Imaging Centre (WCIC) at the IoPPN, King’s College, London, for help with microscopy.

## Supplemental Material

### Supplemental Figures

**Supplemental Figure 1.**
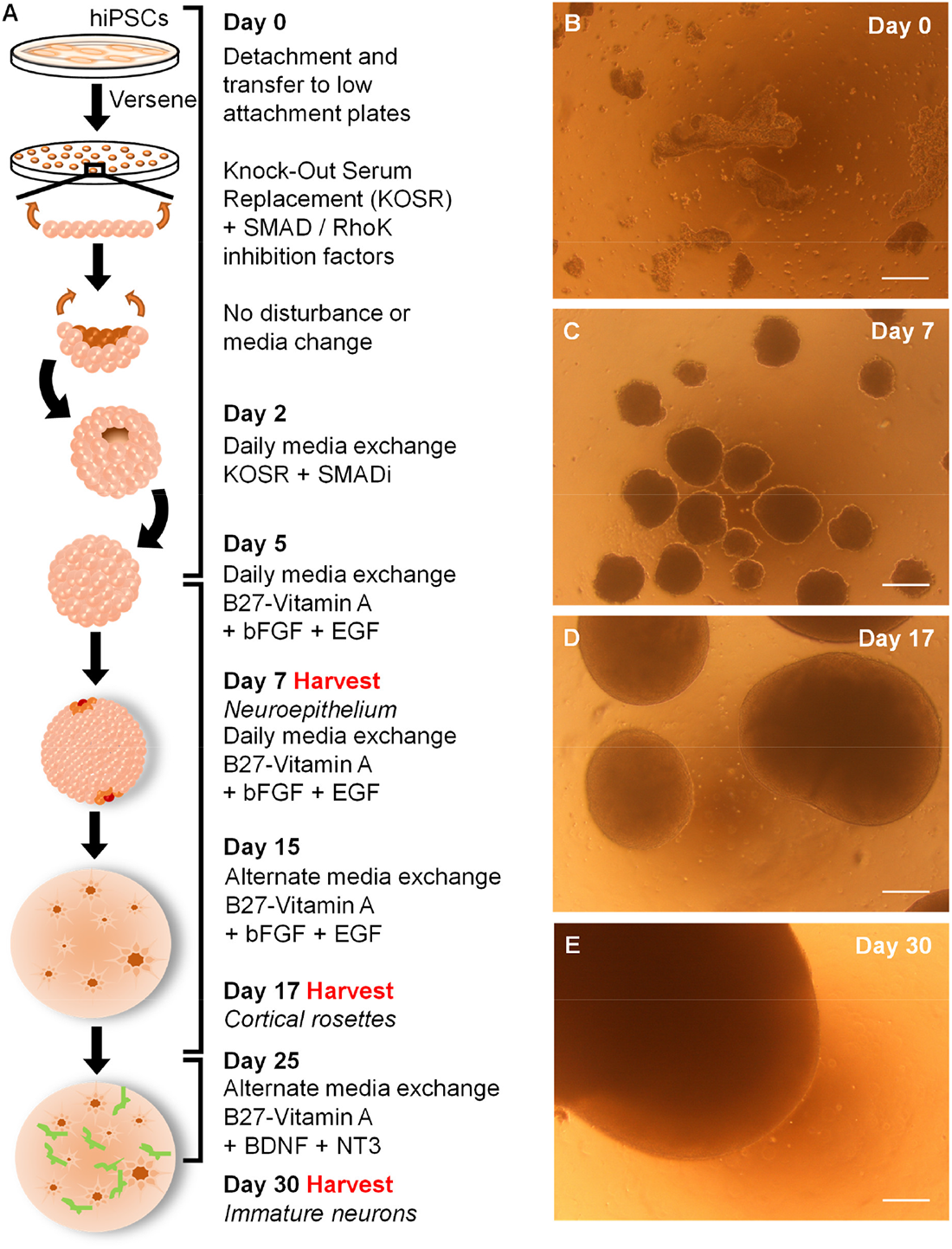
Overview of 3D cortical spheroid (hCS) generation. **(A)** Schematic describing methodology used for 3D neuralization of hiPSCs (adapted from Paşca et al., 2015). **(B-E)** Bright field micrographs of hiPSCs undergoing 3D neuralization. **(B)** Day 0, confluent hiPSC colonies are detached as intact sheets and transferred to suspension culture to allow 3D morphogenesis alongside dual SMAD inhibition. **(C)** Day 7, an early neuroepithelial fate is achieved and roughly spherical structures form. HCS at day were ~100-300 μm in diameter. **(D)** Day 17, rapid progenitor expansion results in large neuroepithelia formation, with smooth radial organisation at edges as expected during corticogenesis, indicated by white arrows. At this time point, hCS were ~1 mm in diameter. **(E)** Day 30, cortical spheroids approach maximum size, often >2 mm in diameter, with growth factors supporting differentiation to post-mitotic immature neurons. Scale bar = 250 μm.

**Supplemental Figure 2.**
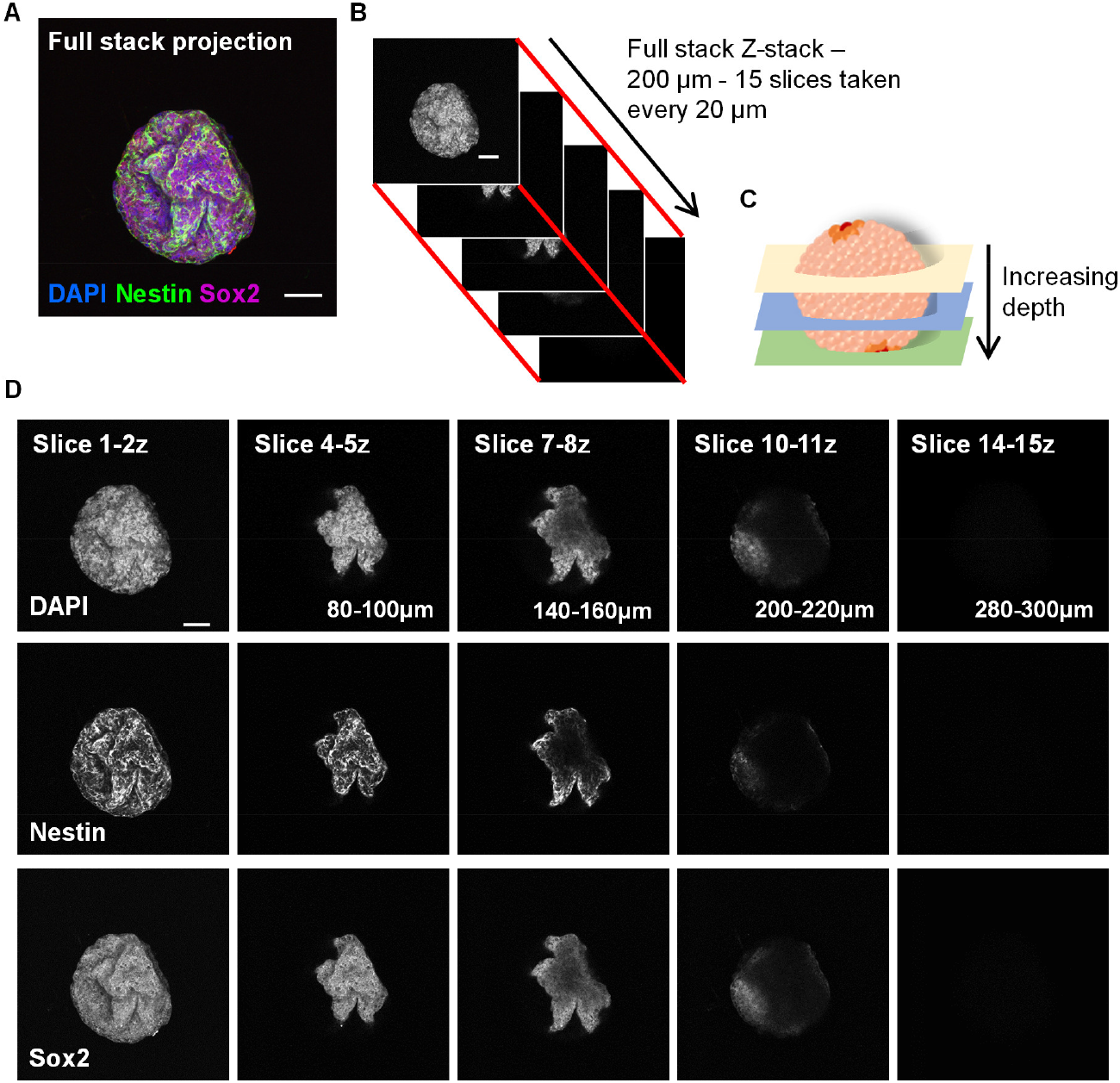
Confocal images of day 7 hCS at increasing depths. **(A)** Composite maximal intensity projection of day 7 hCS shown in Figure 2. **(B)** Visual representation of confocal images that comprise z-stack. **(C)** Schematic of how images within z-stack correspond to overall hCS structure. **(D)** Confocal images of each imaged channel taken at differing depths (axial plane). A significant drop off in intensity is seen after 100 μm. Scale bar = 50 μm.

**Supplemental Figure 3.**
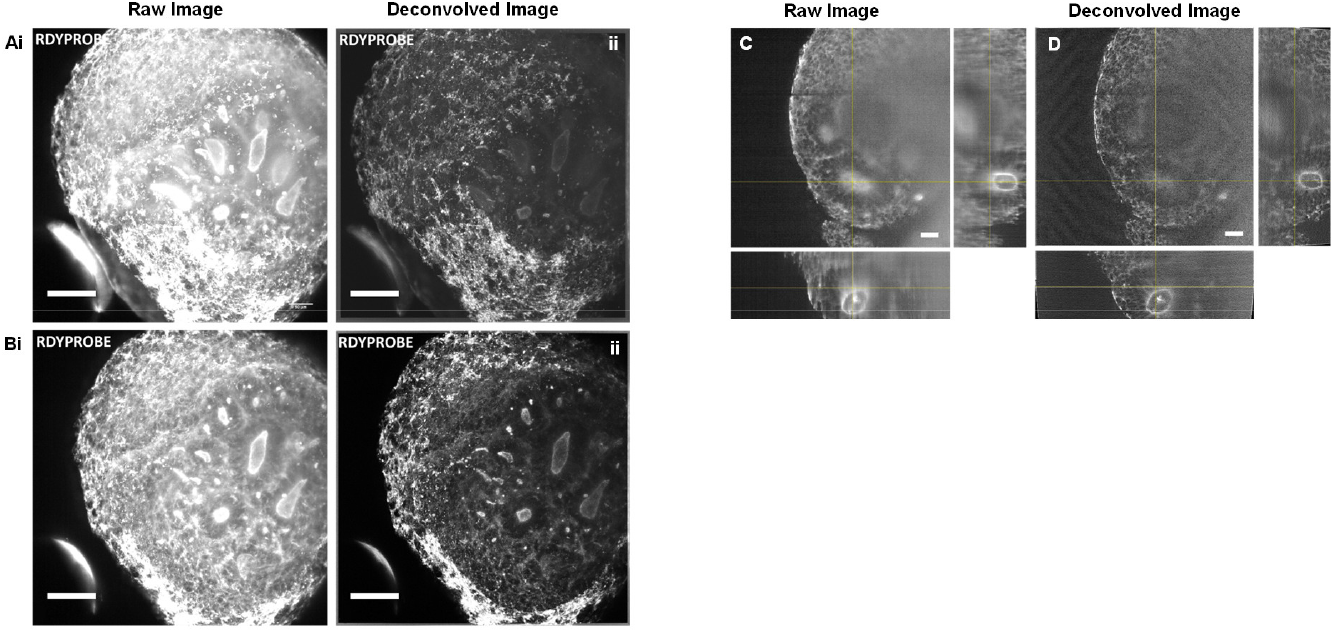
Comparison of raw and deconvolved light sheet acquisitions of hCSs. **(A)** A raw **i)** and deconvolved **ii)** output max intensity Z stack, showing the full stack (500z, 200 μm) from acquisition including the outer edges of tissue. **(B)** A raw **i)** and deconvolved **ii)** section of the previous Z stack A) (60z-360z), removing the outer and deep edges of tissue permitting unobstructed visualisation of internal structures. Scale bars, 100 μm. Phallodin-488 Readyprobe (F-actin) on Day 30 hCSs. **(C)** An orthogonal view of a raw output Z stack image displaying cortical rosettes. **(D)** An orthogonal view of the same Z stack image as **(C)** but deconvolved. Scale bars = 50 μm.

**Supplemental Figure 4.**
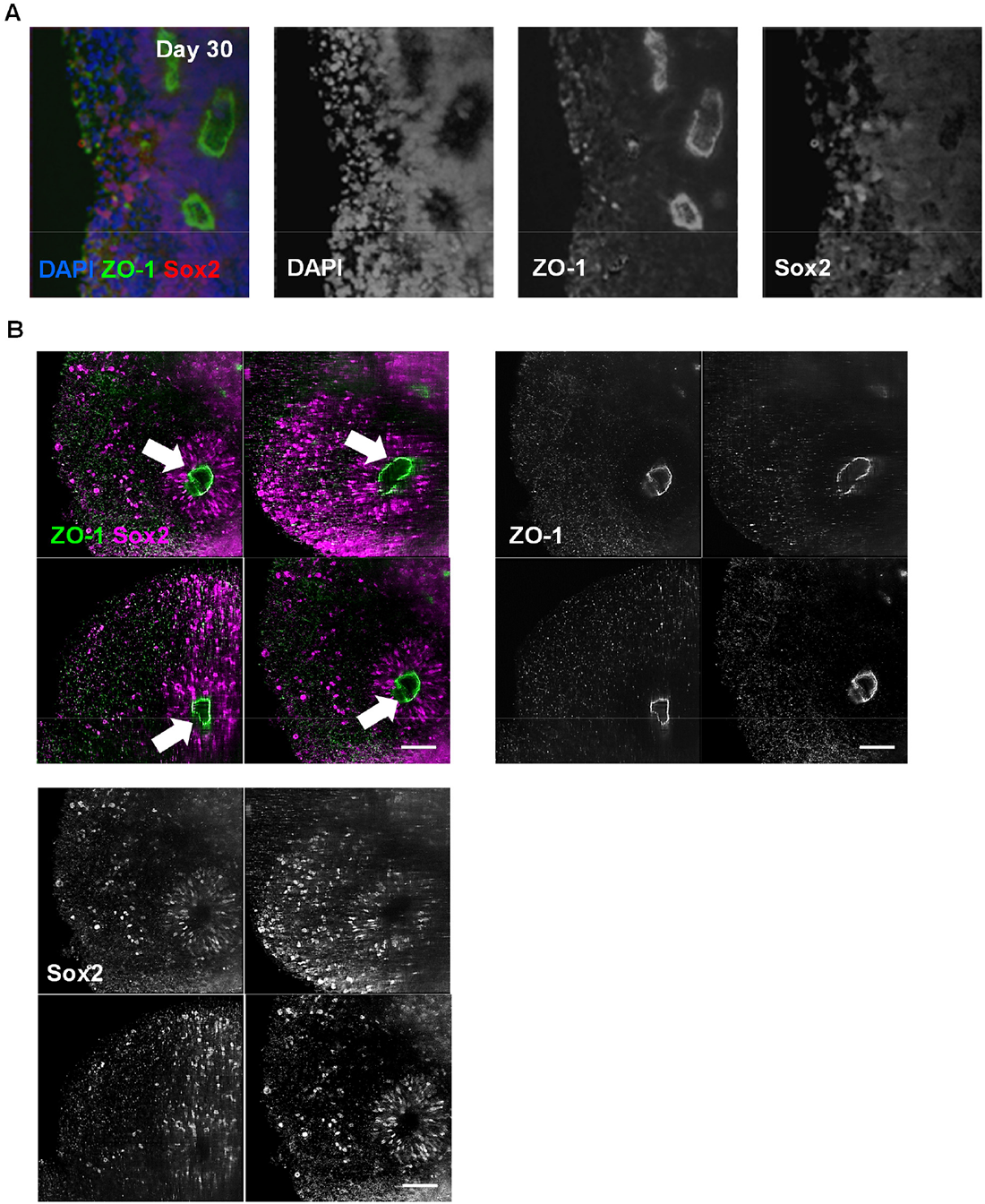
ALSM imaging of cleared day 30 hCS. **(A)** Maximum intensity Z stack (2D) projections (composite and individual channels), showing the full stack of cleared day 30 hCS immunostained for ZO-1 (green), Sox2 (red) and DAPI (blue). **(B)** Maximum intensity Z stack (2D) projections of optical sections shown in main Figure 5 C. Images are composite for ZO-1 (green) and Sox2 (magenta) channels, and subsequently of ZO-1 and Sox2 channels individually as grey scale images. Scale bars = 100 μm.

**Supplemental Figure 5.**
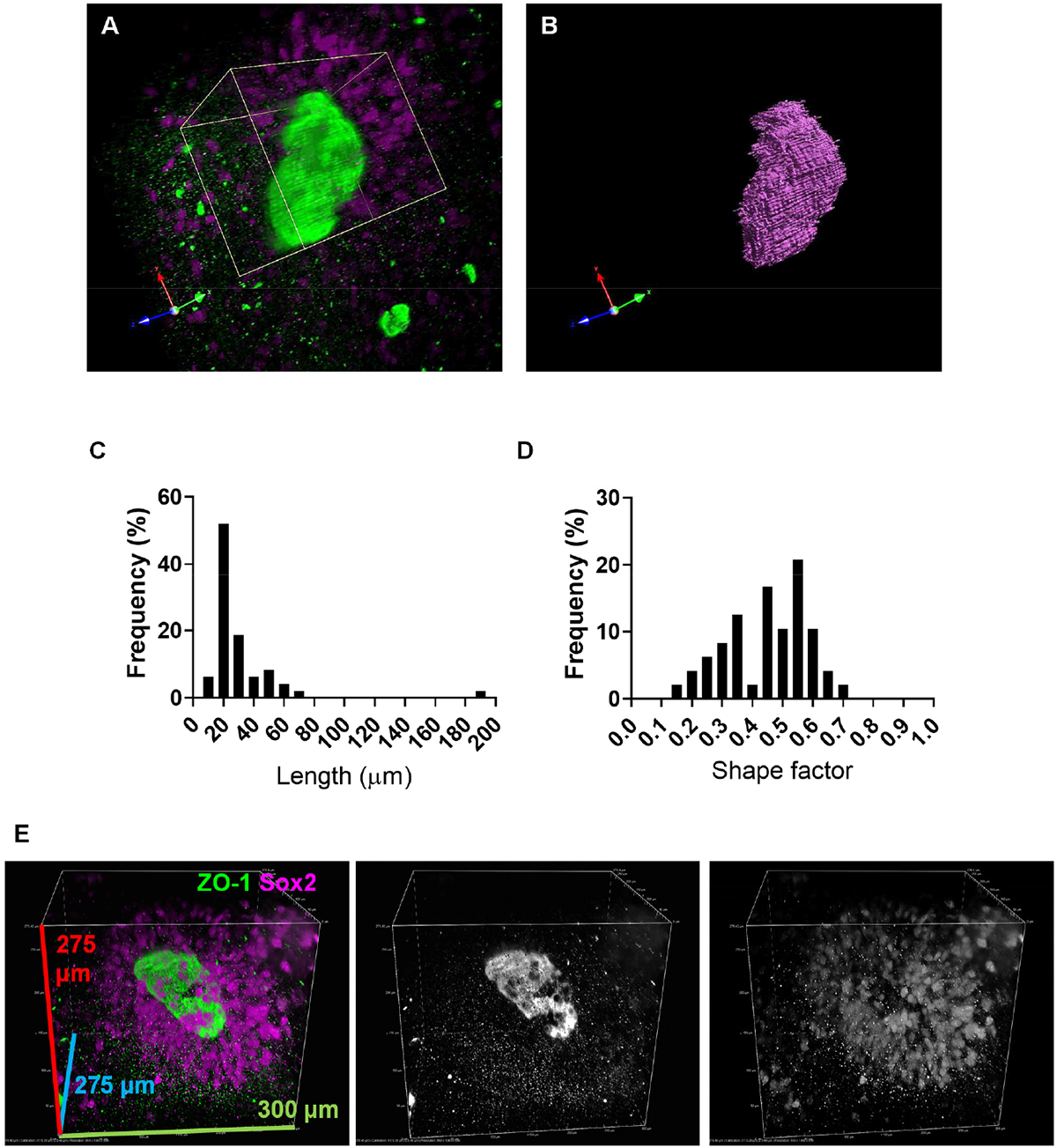
ALSM imaging of neural rosette lumens in cleared day 30 hCS. **(A)** Alternative digital zoom of largest neural rosette lumen shown in main Figure 6E. Green channel indicates ZO-1-psotive apical membrane; magenta channel indicates Sox2-posiive NPCs surrounding central lumen. **(B)** Threshold image of ZO-1-positve lumen demonstrating morphology of structure in 3D. **(C & D)** Frequency distribution of neural rosette lumen length (μm; **C**) and shape factor (**D**). **(E)** Alternative representative 3D render of largest neural rosette lumen (shown in main figure 6G) demonstrating the radial organisation of Sox2-positve NPCs surrounding ZO-1-positive (green) rosette lumen.

**Supplemental Figure 6.**
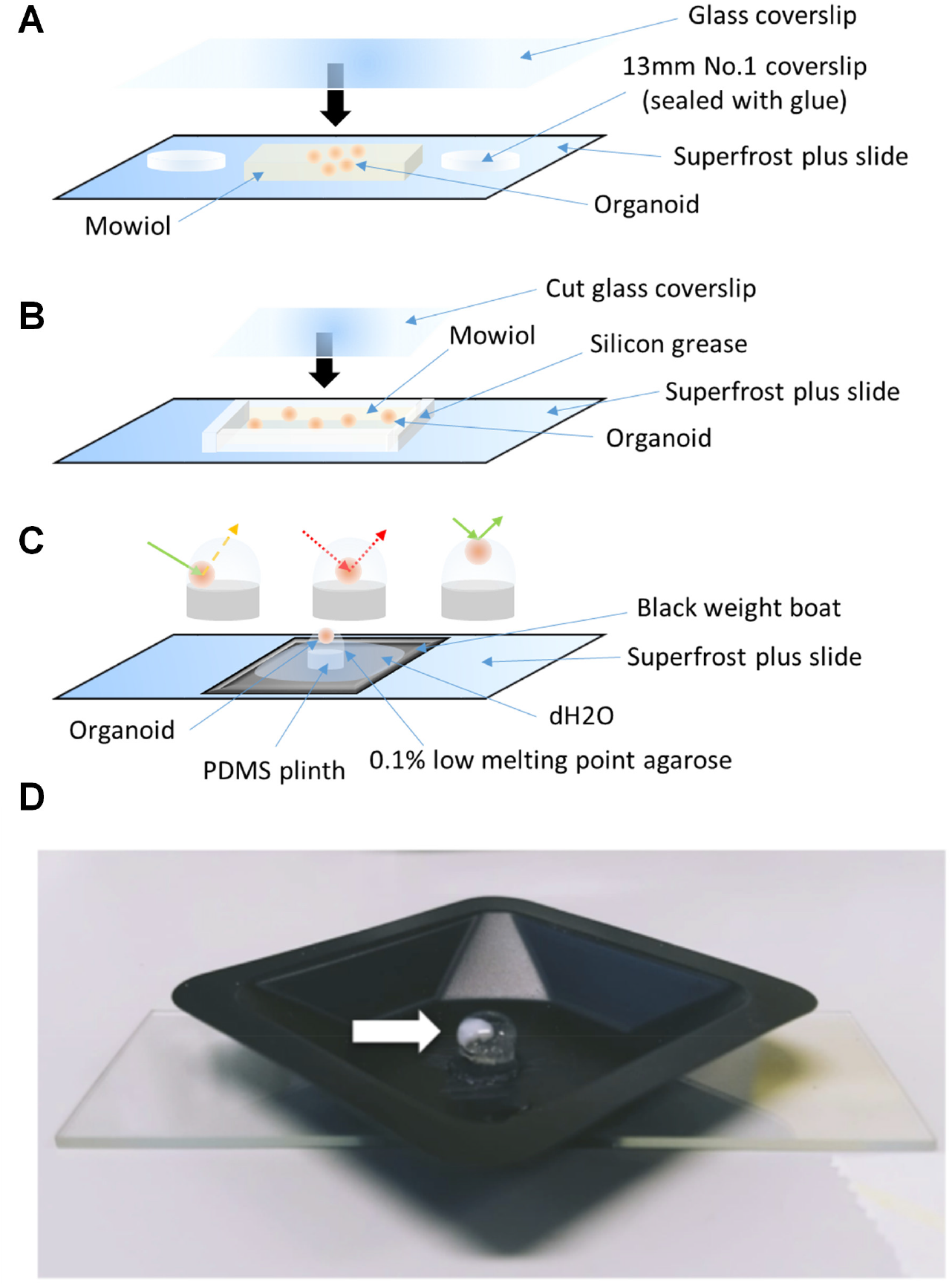
Preparations of setups used for mounting intact 3D tissues. Figure displaying the mounting setups developed for imaging of spheroids by confocal and ALSM. **(A)** The prototype setup used for initially imaging Day 7 hCS using a confocal microscope. Despite sealing the spheroids between the glass, Mowiol was free to exit the setup if not laid flat upon storage, in addition spheroids were larger than initially predicted (~300μm) thus may have been distorted as No.1 coverslips are only 100μm thick. **(B)** An improved design of **(A)** that uses a layer of silicon grease surrounding the spheroids to buffet incoming pressure. However, this method does not offer the same level of protection as hardened glass and ultimately Mowiol solution can leak if pressure is too intense. **(C)** A setup designed by M Squared Life for use with their ALSM. A continuous film of water must cover both the agarose and the objectives used for imaging to reduce visual aberrations. In addition, it is of utmost importance to place the sample tissue as close to the peak of the agarose hemisphere as possible, as to prevent the distance that laser focused light must travel before being refracted into the emitter, and to ensure the lowest amount of light scatter to reduce background signal. **(D)** A second iteration of the imaging setup for use with ALSM. This setup was also used for clearing of hCSs prior to immunostaining and subsequent imaging.

### Supplemental Movie

**Supplemental Movie 1.** Three-dimensional ‘fly-through’ of ALSM volumetric image of cleared day 30 hCS immunostained for NPC marker Sox2 (magenta) and aplical membrane marker ZO-1 (green).

**Supplemental Movie 2.** Three-dimensional ‘fly-through’ digitally zoomed onto largest neural rosette lumen of cleared day 30 hCS. HCS were immunostained for NPC marker Sox2 (magenta) and aplical membrane marker ZO-1 (green).

### Supplemental Tables

**Supplemental Table 1:** Primary antibodies used in study.

| **Primary Antibody** | **Species** | **Supplier** | **Cat. No.** |
| --- | --- | --- | --- |
| Pax6 | Rb | BioLegend | 901301 |
| Tuj1 | Ms | BioLegend | 801201 |
| N-Cad | Ms | Sigma Aldrich | C3865 |
| ZO-1 | Rb | Thermo Fisher | 40-2200 |
| PKCλ | Ms | BD Bioscience | 610207 |
| DCX | Rb | Abcam | ab18723 |
| MAP2 | Chk | Abcam | ab92434 |
| Nestin | Ms | Merck Millipore | MAB5326 |
| Sox2 | Rb | Merck Millipore | 2003600 |

**Supplemental Table 2:** ScaleS solution components used in study.

| **Ingredients** | **ScaleSQ(5)** | **ScaleS4(0)** |
| --- | --- | --- |
| D-(–)-sorbitol (w/v)% | 22.5 | 40 |
| Glycerol (w/v)% | - | 10 |
| Urea (M) | 9.1 | 4 |
| Triton X-100 (w/v)% | 5 | - |
| Dimethylsulfoxide (v/v)% | - | 25 |
| pH | 8.2 | 8.1 |

